# A UDP-GlcNAc : βGal β1-6 GlcNAc transferase involved in bloodstream form *N*-glycan and procyclic form GPI anchor elaboration in *Trypanosoma brucei*

**DOI:** 10.1101/2021.06.21.449264

**Authors:** Samuel M. Duncan, Rupa Nagar, Manuela Damerow, Dmitry V. Yashunsky, Benedetta Buzzi, Andrei V. Nikolaev, Michael A.J. Ferguson

## Abstract

*Trypanosoma brucei* has large carbohydrate extensions on its *N*-linked glycans and glycosylphosphatidylinositol (GPI) anchors in its bloodstream form (BSF) and procyclic form (PCF), respectively. The parasite’s glycoconjugate repertoire suggests at least 38 glycosyltransferase (GT) activities, 16 of which are unknown. Here, we probe the function(s) of a putative β3GT gene, *TbGT10*. The BSF null mutant is viable *in vitro* and *in vivo* and can differentiate into PCF, demonstrating non-essentiality. However, the absence of TbGT10 led to impaired elaboration of *N*-glycans and GPI anchor sidechains in BSF and PCF parasites, respectively. Glycosylation defects include reduced BSF glycoprotein binding to ricin and to monoclonal antibodies mAb139 and mAbCB1. The latter bind a carbohydrate epitope of lysosomal glycoprotein p67 that we show here, using synthetic glycans, consists of (−<u>6</u>Galβ1-4GlcNAcβ1-)_≥ 4_ poly-*N*-acetyllactosamine repeats. Methylation linkage analysis of Pronase glycopeptides isolated from BSF wild-type and TbGT10 null parasites show a reduction in 6-*O*-substituted- and 3,6-di-*O*-substituted-Gal residues. Together, these data suggest that *TbGT10* encodes a UDP-GlcNAc : βGal β1-6 GlcNAc-transferase active in both BSF and PCF life-cycle stages elaborating complex *N*-glycans and GPI sidechains, respectively. The β1-6 specificity of this β3GT gene product and its dual roles in *N*-glycan and GPI glycan elaboration are notable.

## Introduction

The African trypanosomes are protozoan parasites that cause Nagana in cattle and Sleeping Sickness in humans. Infection is maintained by the proliferative “slender form” of the parasite which resides chiefly in the bloodstream and lymphatics of the mammalian host, but also in sub-cutaneous (1) and adipose tissue reservoirs (2). Slender forms survive in their mammalian host by expressing a dense coat of 5 million variant surface glycoprotein (VSG) homodimers (3) tethered to the membrane by glycosylphosphatidylinositol (GPI) anchors (4). VSGs are classified on amino acid sequence motifs, but share general glycosylation features such as a GPI-moiety elaborated by 0-6 galactose residues and the attachment of 1-3 asparagine-linked glycans (*N-*glycans) (4–6). The VSG coat shields the surface membrane from macromolecules such as complement components, whilst enabling the diffusion of small nutrient molecules for uptake into the cell via underlying transmembrane transporters. The *N-* glycosylation of VSG insulates its protein core from intermolecular interactions with proximal surface proteins, enabling dense packing to occur at a level approaching the molecular crowding threshold at which point membrane diffusion is affected (7). VSGs ultimately do not protect from the adaptive immune response as they are immunogenic. Consequently the parasite has evolved a process termed antigenic variation whereby it switches expression to alternative VSGs from a large repertoire of silent genes to maintain infection (3, 8). Glycosylation of VSG can also contribute to the evasion of adaptive immunity, as demonstrated by the discovery that O-glycosylation by 1-3 hexose residues at the top of certain VSGs prolongs infection in mice through antigen heterogeneity (9). Bloodstream form (BSF) *T. brucei* produce a range of *N-*glycans from short Man_3_GlcNAc_2_ paucimannose structures to highly branched, complex and polydisperse poly-*N-*acetyl-lactosamine (poly-LacNAc) including a very large family of structures composed of on average of 54 LacNAc repeats (4, 10–13). Glycoproteins, such as p67, bearing these poly-LacNAc structures localise to the flagellar pocket and the lysosomal/endosomal system (12, 14–16). Poly-LacNAc glycans have been suggested to play a role in endocytosis (17).

Following uptake of transmissible stumpy form BSF parasites in a bloodmeal, differentiation to the procyclic form (PCF) trypanosome occurs in the tsetse fly midgut. During this transformation the VSG coat is replaced by different GPI anchored glycoproteins called procyclins (18). These are characterised by rod-like polyanionic dipeptide (EP) or pentapeptide (GPEET) repeats with and without a single triantennary Man_5_GlcNAc_2_ *N-*linked glycan (19) and without and with threonine phosphorylation (20), respectively. Both types of procyclin share the largest and most complex GPI-anchor sidechains described thus far. These are composed of branched *N-*acetyl-lactosamine (LacNAc; Galβ1-4GlcNAc) and lacto-*N-*biose (LNB; Galβ1-3GlcNAc) containing structures terminating in β-Gal (19, 21) that can be further modified by α2-3-linked sialic acid residues by the action of cell-surface trans-sialidase (21, 22). Surface sialylation of PCF trypanosomes plays an important role in efficient tsetse fly colonisation (23) and the procyclins appear to shield susceptible surface proteins from proteolytic attack the tsetse gut (24).

In summary, BSF GPI anchors have relatively simple Gal-containing sidechains, whereas PCF GPI anchors can be extremely large and complex. The reverse is true for protein *N-*glycosylation where BSF parasites can express extremely large and complex structures, whereas wild type PCF parasites express predominantly oligomannose structures. The control of *N-*glycan type is primarily dictated by oligosaccharyltransferase expression whereby the formation of complex *N-*glycan structures requires the transfer of biantennary Man_5_GlcNAc_2_ from Man_5_GlcNAc_2_ -PP-dolichol by the TbSTT3A oligosaccharyltransferase, whereas oligomannose structures originate from triantennary Man_9_GlcNAc_2_ transferred from Man_9_GlcNAc_2_ -PP-dolichol by the TbSTT3B oligosaccharyltransferase (25–28). Thus, downregulation of TbSTT3A expression in PCF cells switches them away from complex *N-*glycan to oligomannose *N-*glycan expression.

Glycosyl linkages catalysed by glycosyltransferases (GTs) are defined by the configuration (pyranose or furanose) and anomericity (α or β) of the transferred sugar, by the inter-sugar linkage to the aglycone acceptor (e.g. 1-2, 1-3, 1-4 or 1-6 to a hexapyranose sugar acceptor) and by the precise structure of the aglycone acceptor. In general, each different glycosyl linkage is catalysed by a unique GT or family of GTs using a specific activated sugar glycosyl donor, generally a nucleotide sugar or a lipid-linked sugar phosphate. Around 38 unique GTs/GT families are predicted to occur in *T. brucei* to account for the 38 known glycosidic linkages made by the parasite (Supplementary Fig. S1 and Fig. S2). Of these 38 predicted GTs/GT families, 11 have been experimentally identified, 11 predicted by sequence homology and 16 remain to be identified bioinformatically and/or experimentally (Supplementary Table S1).

The GTs that elaborate GPI-anchors and *N-*linked glycans with α-Gal, β-Gal and β-GlcNAc residues are presumed to be UDP-Gal and UDP-GlcNAc dependent. In the genome of *T. brucei*, we can find 19 putative UDP-Gal or UDP-GlcNAc-dependent β-GT sequences (Supplementary Fig. S1 and Table S1), excluding predicated pseudogenes. These belong to a distinct GT family (GT67) in the Carbohydrate Active enZYmes (CAZY) database (29). These genes appear to have evolved from an ancestral inverting glycosyltransferase, similar to the mammalian β3GT family (13, 30). Two of these have been shown to be involved in the synthesis of GPI sidechains in PCF cells: A UDP-Gal : βGlcNAc β1-3 Gal-transferase (TbGT3) (31) and a UDP-GlcNAc : βGal β1-3 GlcNAc transferase (TbGT8) (13). Ablation of either enzyme results in reduced molecular weight procyclins due to aberrant glycosylation of the GPI anchor. However, neither enzyme is essential for growth *in vitro* or for the establishment of tsetse fly infections. Analysis of *N-*linked glycosylation in BSF mutants revealed that TbGT8 deficient mutants also have pronounced changes in *N-*glycosylation, as manifested by reduced wheat germ agglutinin (WGA) and ricin binding (13, 32). The presence of GlcNAcβ1-3Gal inter-LacNAc linkages in both BSF *N-* linked glycans and PCF GPI anchors presumably explains the dual function of TbGT8 and it is possible that other functional dualities may exist for other *Trypanosoma* GTs.

Two other putative UDP-Gal or UDP-GlcNAc-dependent GT sequences, TbGT11 and TbGT15, have been studied and shown to be the equivalents of animal GlcNAc-transferase I (GNTI) and GlcNAc-transferase II (GnTII), respectively (33, 34). These Golgi apparatus enzymes are both UDP-GlcNAc : αMan β1-2 GlcNAc transferases that prime the elaboration of the α1-3 and α1-6 arms of the common Man_3_GlcNAc_2_ core of complex *N-*glycans in BSF parasites. It is notable that, despite sharing a common β3GT ancestor, TbGT11 and 15 both catalyse β1-<u>2</u> glycosidic linkages. We therefore anticipate that members of the *T. brucei* “β3GT” family may also catalyse some or all of the β1-4 and β1-6 glycosyl linkages described in (Supplementary Fig. S1 and S2 and Supplementary Table S1). Neither TbGnTI (33) nor TbGnTII (34) is essential for BSF survival *in vitro* or *in vivo* and there evidence that the absence of elaboration on either arm of the Man_3_GlcNAc_2_ core is compensated for, at least in part, by greater elaboration of the remaining arm.

In this study, we characterised TbGT10 (Tb927.5.2760) a putative GT with a suggested role in the mode of action of the trypanocidal drug suramin (35).

## Results

### Analysis of the TbGT10 predicted amino acid sequence

Tb927.5.2760 encodes a 384-amino acid protein with a theoretical mass of 43.5 kDa. Sequence analysis predicts type II membrane protein topology, common among Golgi resident GTs, with a 12 residue *N-*terminal cytoplasmic domain, a 20 amino acid transmembrane domain and a large Golgi luminal domain. The latter includes a putative galactosyltransferase catalytic domain between residues 180-242 and characteristic GT DXD motif, consistent with coordination to a nucleotide sugar donor via a divalent cation (36).

### Creation and growth phenotype of a bloodstream form TbGT10 conditional *null* mutant

The knockdown of TbGT10 transcripts by tetracycline-inducible RNAi caused a growth defect (Supplementary Fig. S3A), suggesting that TbGT10 might be essential. However, this proved no to be the case, as described below. To determine the function of *TbGT10*, we decided to make a conditional null mutant of this gene. Although we were able to construct Δ*gt10::PAC/GT10* (*GT10*^*Ti*^) clones, we were unable to replace the second *TbGT10* allele in the presence of tetracycline. We therefore decided to take a different approach by applying rapamycin-induced diCre mediated gene deletion (37, 38). To this end, the first allele of *TbGT10* was replaced by a puromycin acetyltransferase (*PAC*) resistance cassette to generate a heterozygote and the remaining allele was replaced with an expression construct containing the *TbGT10* gene flanked by *loxP* sites (floxed), in tandem with a hygromycin phosphotransferase-thymidylate kinase (*HYG-TK*) cassette for negative and positive selection, respectively (39). A single Δ*gt10*::*PAC*/Δ*gt10*::*TbGT10-HYG-TK*^*Flox*^ clone was selected, expressing only floxed *TbGT10*. Finally, a modified version of pLEW100v5 lacking the Tet operator was used to constitutively express both diCre subunits under phleomycin (PHLr) selection at the ribosomal small subunit (*SSU*) locus (Fig. 1A). Δ*gt10*::*PAC*/Δ*gt10*::*TbGT10-HYG-TK*^*Flox*^ (*SSU diCre*) clones (hereafter referred to as *TbGT10* conditional knockout (cKO) cells) were assayed for floxed *TbGT10-HYG-TK* locus excision in the presence of 100 nM rapamycin by PCR analysis (Fig. 1B), and for loss of hygromycin resistance (Fig. 1C). A *TbGT10* cKO clone showing these characteristics was selected for further analysis.

**Figure 1.**
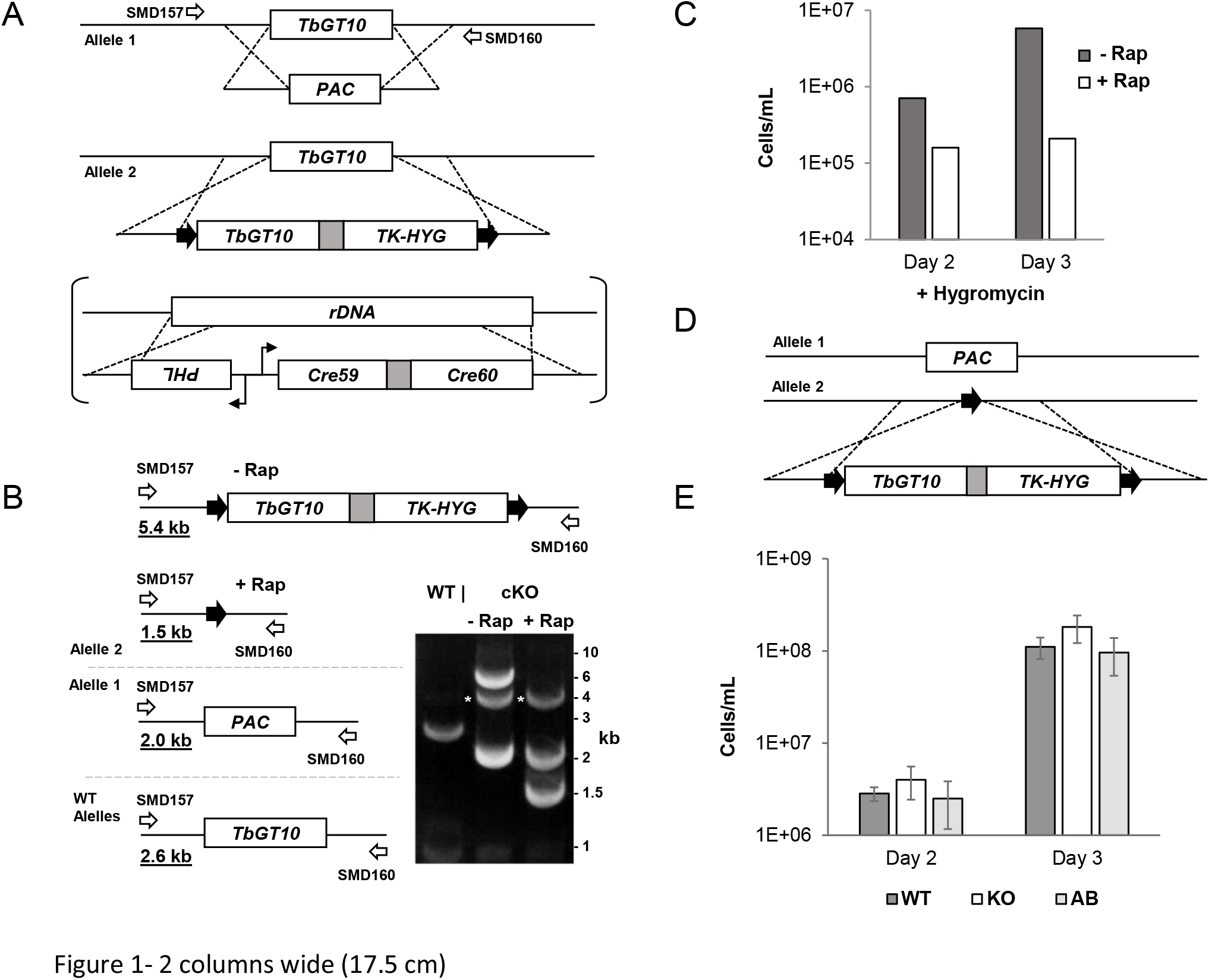
Generation of bloodstream form *TbGT10* conditional null mutant. A. Gene replacement strategy to generate *TbGT10* cKO cells by replacement of one *TbGT10* allele with a *PAC* gene, introduction of a *loxP* (black arrows) flanked TbGT10 transgene and subsequent insertion of a constitutive diCre expression cassette at the ribosomal DNA locus (in squared brackets). B. Gene excision was analysed by PCR amplification. Schematic shows the *TbGT10*^*Flox*^ locus and the recombination event after rapaymycin treatment. PCR amplification of gDNA harvested 48 h after rapamycin treatment was performed using oligonucleotide primers SMD157 and 160 (open arrows) that anneal outside of the homologous recombination site. Expected amplicon sizes are underlined. Resolution of PCR products by agarose gel electrophoresis confirms the replacement of endogenous *TbGT10* by *TbGT10*^*Flox*^ and the excision of *TbGT10*^*Flox*^ upon rapamycin treatment. A non-specific 4 kb amplicon was observed following PCR amplification with *TbGT10* cKO gDNA as template (white asterisks). C. Growth of *TbGT10* cKO cells cultured in the presence (+ Rap) or absence (−Rap) of 100 nM rapamycin for 3 days followed by seeding in the presence or absence of hygromycin to assess floxed gene loss by hygromycin sensitivity. D. Strategy for generation of *TbGT10* add-back (AB) mutants by re-introduction of *TbGT10*^*Flox*^ at the excised locus of *TbGT10* KO mutants. D and E. Infectivity of wild-type (WT), *TbGT10* KO and *TbGT10* AB mutant cells in mice. Mice were infected with 2 × 10^5^ cells and the number of cells per ml of blood was counted 2 and 3 days post-infection. No significant difference in infectivity was observed.

Although rapamycin treatment has previously been suggested to be toxic to BSF *T. brucei* (40), we find that that 100 nM rapamycin is well tolerated by BSF *T. brucei* and that, in our hands, rapamycin has an EC50 concentration of 5.9 µM (Supplementary Fig. S3B).

Upon rapamycin*-*induced excision of the single *TbGT10* gene in our cKO clone, we observed a slightly reduced growth rate compared with uninduced cells (Supplementary Fig. S3C). To generate a stable null mutant cell line, *TbGT10* cKO cells were induced with 100 nM rapamycin for 5 days and a *TbGT10* knockout (KO) clone was selected by serial dilution. The absence of *TbGT10* in the *TbGT10* KO clone was confirmed by PCR (Supplementary Fig. S3D) and this clone exhibited normal growth kinetics *in vitro*, suggesting some adaptation had taken place to recover a normal (wild type) growth phenotype. We further complemented this *TbGT10* KO clone by re-introducing the floxed TbGT10 expression construct at the *loxP* excised locus, generating a TbGT10 re-expressing line, hereafter referred to as *TbGT10* add-back (AB) (Fig. 1D). No difference in the ability of wild-type, *TbGT10* KO or *TbGT10* AB cells to infect Balb/c mice was detected (Fig. 1E), indicating that TbGT10 is not required for infectivity to mice. These data clearly demonstrate, despite the RNAi result (Supplementary Fig. 3A), that *TbGT10* is not an essential gene for BSF *T. brucei in vitro* or *in vivo*.

### TbGT10 is involved in complex *N-*glycan processing in bloodstream form *T. brucei*

Since the loss of TbGT10 did not compromise cell viability in culture, we were able to study the *TbGT10* cKO and *TbGT10* KO cell lines for effects on protein glycosylation. In the first instance, we looked at lectin blotting of SDS cell lysates. Blotting with ricin (*Ricinius communis* agglutinin; RCA) indicated a reduction in terminal β-galactose residues in TbGT10 deficient lysates compared to those of wild-type or *TbGT10* AB cells (Fig. 2; Supplementary Fig. S4). This difference manifested itself in the >75 kDa apparent molecular weight glycoproteins, whereas the RCA-reactivity of a band at ∼50kDa (presumed to be VSG221, which carries a terminal β-Gal residue on its GPI anchor) was unaffected (Fig. 2).

**Figure 2.**
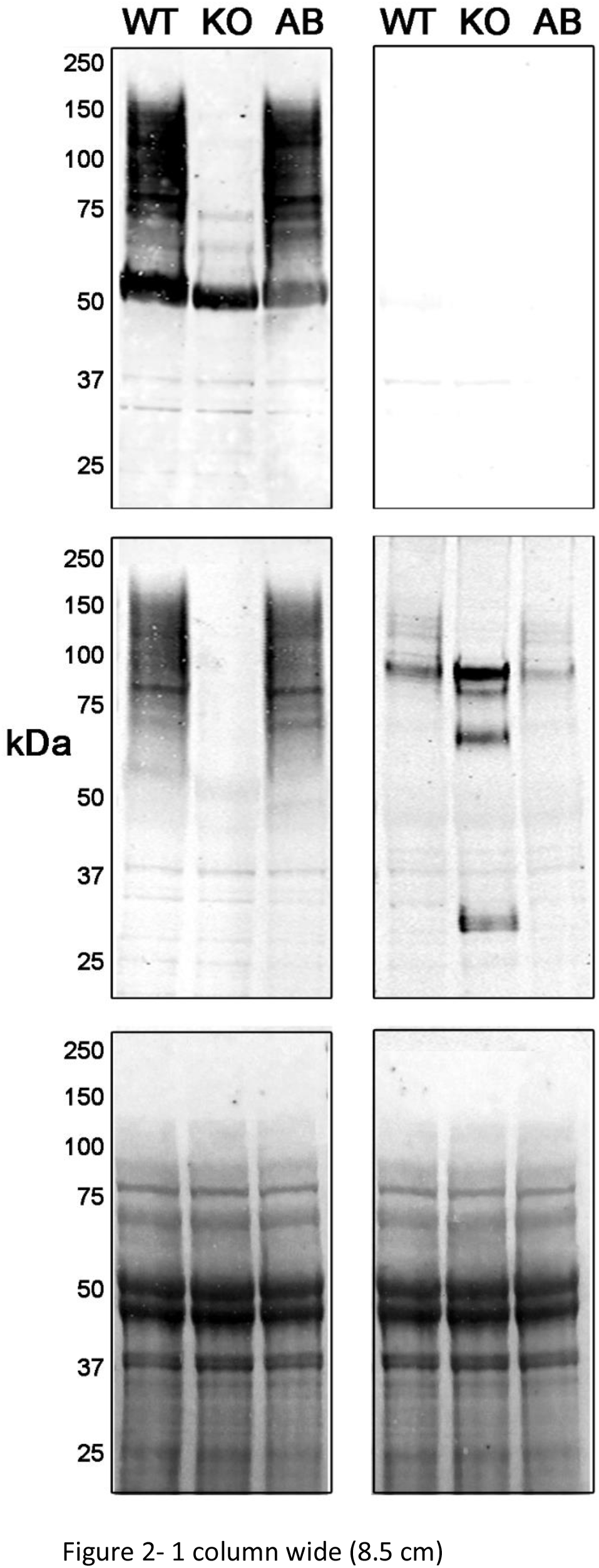
Lectin and Western blotting of *TbGT10* null and add-back glycoproteins. Upper panel, lysates of wild-type (WT), *TbGT10* KO and *TbGT10* AB cells were subjected to SDS-PAGE and transferred to nitrocellulose membrane in duplicate. Membranes were incubated with biotinylated ricin without (left hand side) or with (right hand side) pre-incubation with 10 mg/mL galactose and lactose as a binding specificity control. Middle panel, Western blotting performed with anti-carbohydrate mAb139 (left hand side) and anti-peptide p67C antibodies (right hand side). Lower panel, equal loading and transfer are demonstrated by Ponceau S staining.

To explore the *TbGT10* KO glycosylation phenotype further, we looked at the status of p67, a heavily-glycosylated and proteolytically-processed type I transmembrane glycoprotein carrying up to 14 *N-* linked glycans (14). In BSF cells, nascent p67 in the ER has an apparent MW of 100 kDa which, upon trafficking to the Golgi, becomes processed to ∼150 kDa through the elaboration of some of the *N-* linked glycans to complex poly-LacNAc containing structures. The glycoprotein is then trafficked to the lysosome and cathepsin L-like processing cleaves p67 into a variety of smaller fragments (14). We analysed the fate of p67 in *TbGT10* cKO and *TbGT10* KO cells by probing Western blots of whole cell lysates with monoclonal antibodies mAb139 (an IgG2) (Fig. 2; Supplementary Fig. S4) and mAbCB1 (an IgM) (Supplementary Fig. S5) both known to react with an uncharacterised carbohydrate epitopes or epitopes present on fully processed BSF p67 (41). Immunoreactivity with both antibodies was greatly reduced in lysates of *TbGT10* cKO cells treated with rapamycin compared with untreated controls (Supplementary Fig. S4 and S5) and mAb139 detection was completely ablated in the *TbGT10* KO cells (Fig. 2). These results indicate that the carbohydrate epitope(s) identified by them is/are dependent on TbGT10 activity. Outgrowth of *TbGT10* cKO cells expressing the mAb139 epitope occurred when cells treated with 100 nM rapamycin for 3 days were then grown in the absence of the ligand for a further 9 days, suggesting that conditional loss of *TbGT10* imparts a fitness cost (Supplementary Fig. S4).

To test whether this loss of carbohydrate epitope signal might be due to a reduction in p67 polypeptide expression, we also probed the blots with an affinity purified anti-p67 peptide polyclonal antibody that recognises the C-terminus of the protein (p67C) (Fig. 2). The data suggest that the absence of TbGT10 actually increases the expression of, or stabilises, p67 in BSF cells. Interestingly, the pattern of p67 polypeptide labelling seen (Fig. 2, KO lane) is highly reminiscent of that described for radiolabelled p67 in PCF cells where conversion from the 100 kDa to 150 kDa form does not occur (14).

Genome wide RIT-Seq analysis of BSF cells previously identified both p67 and TbGT10 as contributing to the mode of action of suramin, with subsequent RNAi of p67 leading to 2.6-fold resistance to suramin (35). We investigated the effect of TbGT10 on suramin sensitivity using our TbGT10 cKO cell line (Supplementary Fig. S3E). Rapamycin-induced excision of *TbGT10* conferred a 2.8-fold resistance to suramin compared with uninduced cells. Since RNAi of p67 in BSF cells results in aberrant lysosomal turnover or export (15, 16), the accumulation of processed peptides in *TbGT10* cKO cells (Fig. 2, KO lane) may be a consequence of impaired p67 function, leading to increased suramin resistance.

Taken together, these data suggest that removal of TbGT10 leads to impaired glycosylation and proteolytic processing of BSF p67. However, these aberrations do not appear to affect BSF cell viability *in vitro* or *in vivo*.

### Characterisation of the mAb139 and mAbCB1 epitopes suggests TbGT10 is involved in the synthesis of β1-6 inter-LacNAc linkages

The effects of *TbGT10* excision on mAb139 and mAbCB1 immunoreactivity suggested that definition of the epitope(s) recognised by these monoclonal antibodies could be key to understanding TbGT10 activity. Previous experiments have shown that the immunoreactivity of mAbCB1 to BSF cell lysates is ablated by PNGaseF treatment and by pre-incubation of mAbCB1 with 0.2 M lactose or 0.2 M D-galactose (41). These data suggested to us that the mAbCB1 and mAb139 epitope(s) might be present on the novel poly-LacNAc-containing *N-*glycans described in (12). Within those structures, we considered poly-LacNAc structures with β1-<u>6</u>inter-LacNAc repeat glycosidic linkages to be the most likely epitopes, as these are different from the poly-LacNAc structures with β1-<u>3</u> inter-LacNAc repeat glycosidic linkages that are predominant (i.e., self) in mammalian glycoconjugates. To test this hypothesis, we performed bio-layer interferometry (BLI) using a set of synthetic saccharides (42) and (Supplementary Fig. 6) each conjugated to biotin and containing two to five Galβ1–4GlcNAc (LacNAc) repeats with β1-6 inter-LacNAc repeat glycosidic linkages (Fig. 3). These structures were bound to avidin*-*coated sensor pins and used to detect the binding of mAb139. The data corroborated the hypothesis and indicated at least three β1-6 interlinked LacNAc repeats are required for detectable mAb139 binding, with ≥ 4 LacNAc repeats being optimal. Similar data were obtained using the mAbCB1 antibody (Supplementary Fig. S7), suggesting they have the same epitope specificity.

**Figure 3.**
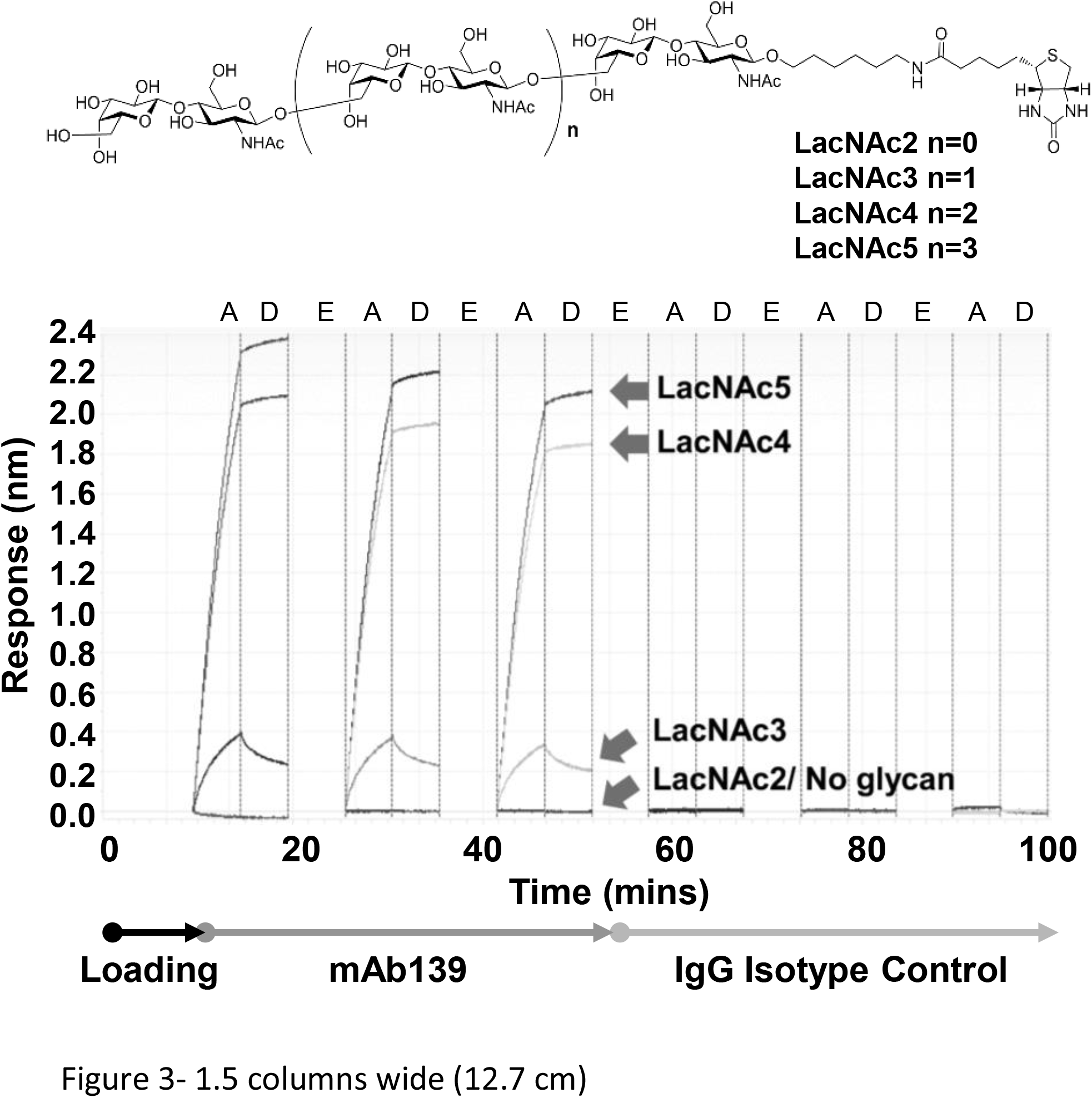
Characterisation of the mAb39 epitope by bio-layer interferometry. The structures of the synthetic biotinylated (−<u>6</u>Galβ1-4GlcNAcβ1-) LacNAc oligosaccharides and their names (LacNAc2-5) are shown at the top. Responses are monitored at the detector, and reported on a sensorgram as a change in wavelength (nm shift) through cycles of association (A), disassociation (D) and elution (E) of mAb139 to and from the LacNAc2-5 loaded sensor pins are shown, alongside sensorgrams using an unrelated IgG2 antibody isotype control.

Based on these data, we suggest that the loss of TbGT10 likely results in the loss of β1-6 interlinked poly-LacNAc repeats in BSF *T. brucei*. Consistent with this, we noted that mAb139 antibody signals in Western blots against whole trypanosome lysates are actually stronger in *TbGT8* null mutants (Supplementary Fig. S8). TbGT8 is a known β1-3 GlcNAc-transferase (13), responsible for catalysing the β1-3 inter-LacNAc glycosidic linkages also found in BSF trypanosome poly-LacNAc *N-*glycans. Thus, we postulate that TbGT10 activity compensates for the absence of TbGT8 activity in those mutants and/or that the presence of GlcNAc β1-6(GlcNAc β1-3)Gal branchpoints reduces mAb139 binding.

### GC-MS methylation linkage analysis of glycoproteins from *TbGT10* null mutants confirms TbGT10 is a β1-6 GlcNAc-transferase

For this study, we developed a new protocol for comparing the carbohydrate structures of wild-type and TbGT mutants, whereby trypanosome ghosts after osmotic shock (i.e., depleted of VSG due to the action of endogenous GPI-specific phospholipase C (43, 44)), are digested with Pronase to yield glycopeptides from all the remaining glycoproteins. The Pronase glycopeptides are freed from peptides and amino acids by diafiltration and from lipidic material by chloroform extraction prior to monosaccharide composition and methylation linkage analysis by GC-MS. The composition analyses indicated a reduction in Gal and GlcNAc relative to Man in the *TbGT10* KO mutant sample compared to the WT sample (Supplementary Fig. S9). The results of the methylation linkage analyses are shown in (Fig. 4 and Table 1). In both samples, we observe similar levels of non*-*reducing terminal- and 2-*O*-substituted-Man residues from oligomannose *N-*glycans and similar levels of 3,6-di-*O*-substituted-Man from oligomannose and complex *N-*glycans. However, the *TbGT10* KO mutant contains less 4-*O*-substituted GlcNAc and significantly less 6-*O*-substituted- and 3,6-di-*O*-substituted-Gal residues than the wild-type. This is consistent with *TbGT10* encoding a UDP-GlcNAc : βGal β1-6 GlcNAc-transferase responsible for the majority of the β1-6 inter LacNAc repeat glycosidic linkages. The increase in 3-*O*-substituted-Gal residues in the *TbGT10* KO sample, compared to wild-type, also suggests that removal of TbGT10 is compensated by increased β1-3 GlcNAc-transferase activity, most likely from TbGT8.

**Figure 4.**
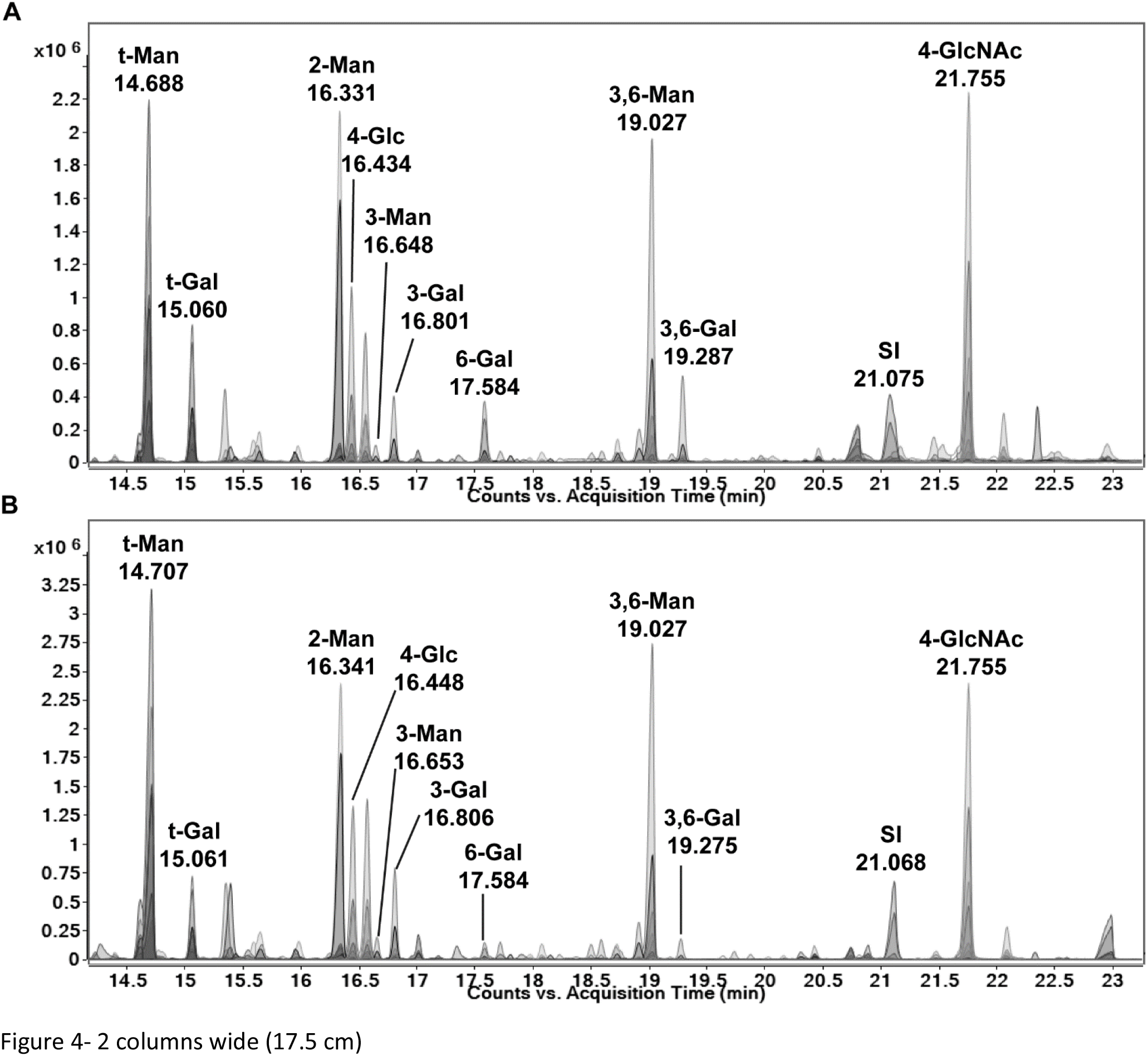
GC-MS methylation linkage analysis of glycoproteins from *TbGT10* null cells. GC–MS linkage analysis of partially methylated alditol acetate (PMAA) derivatives from the Pronase glycopeptide fractions of wild type (upper) and *TbGT10* null mutant (lower) cells. The PMAA peaks are annotated according to the original substitution pattern of the native glycans. For example, t-Gal refers to non-reducing-terminal galactose and 3-Gal refers to 3-O-substituted galactose (see Table 1). The plots are of merged extracted ion profiles of characteristic PMAA fragment ions at *m/z* 102,117, 118, 129, 145, 159, 161, 162, 168, 189, 205, 210, 233 and 234.

**Table 1.** GC-MS methylation linkage analysis of Pronase glycopeptides from wild type and *TbGT10* null mutant cells. The Pronase glycopeptides were permethylated, hydrolyzed, deutero-reduced, and acetylated to yield PMAAs for analysis by GC-MS. Residue types were deduced from the electron-impact mass spectra and retention times.

| PMAA derivatives | Residue types | Retention time(min) | Absolute counts in Wild type (%) | Absolute counts in GT10 Null (%) |
| --- | --- | --- | --- | --- |
| [1-<sup>2</sup>H]-1,5-Di-*O*-acetyl-2,3,4,6-tetra-*O*-methylmannitol | t-Man | 14.68 | 14979665 (100) | 21836774 (100) |
| [1-<sup>2</sup>H)]1,2,5-Tri-*O*-acetyl-2,4,6-tri-*O*-methylmannitol | 2-Man | 16.33 | 13705181<br>(91.49) | 15713041<br>(71.95) |
| [1-<sup>2</sup>H)]-1,3,5-Tri-*O*-acetyl-2,4,6-tri-*O*-methylmannitol | 3-Man | 16.64 | 1283402<br>(8.56) | 1810740<br>(8.29) |
| [1-<sup>2</sup>H)]-1,3,5,6-Tetra-*O*-acetyl-2,4-di-*O*-methylmannitol | 3,6-Man | 19.02 | 9978554<br>(66.61) | 12864144<br>(58.91) |
| [1-<sup>2</sup>H)]-1,5-Di-*O*-acetyl-2,3,4,6-tetra-*O*-methylgalactitol | t-Gal | 15.06 | 7015253<br>(46.83) | 5931412<br>(27.16) |
| [1-<sup>2</sup>H)]-1,3,5-Tri-*O*-acetyl-2,4,6-tri-*O*-methylgalactitol | 3-Gal | 16.80 | 2522793<br>(16.84) | 5137931<br>(23.52) |
| [1-<sup>2</sup>H)]-1,5,6-Tri-*O*-acetyl-2,3,4-tri-*O*-methylgalactitol | 6-Gal | 17.58 | 3312478<br>(22.11) | 1571471<br>(7.19) |
| [1-<sup>2</sup>H)]-1,3,5,6-Tetra-*O*-acetyl-2,4-di-*O*-methylgalactitol | 3,6-Gal | 19.28 | 3817044<br>(25.48) | 1512096<br>(6.92) |
| [1-<sup>2</sup>H)]-1,4,5-Tri-*O*-acetyl-2-methylacetamido-3,6-di-*O*-methylglucosaminitol | 4-GlcNAc | 21.75 | 11448507<br>(76.42) | 9976630<br>(45.68) |

### TbGT10 also elaborates procyclin GPI sidechains

Since the deletion of the TbGT8 β1-3 GlcNAc-transferase activity is known to affect both BSF *N-*glycans and PCF GPI sidechains in *T. brucei* (13), we also sought to analyse the procyclins of the *TbGT10* KO mutant. Using methodology described previously (13), BSF WT, *TbGT10* KO and *TbGT10* add-back cells were differentiated to PCF cells *in vitro* by culturing them in SDM79 medium containing 3mM citrate and *cis*-aconitate at 28°C. Procyclins were harvested from these cells 4 days after differentiation by differential solvent extraction, resolved by SDS-PAGE gel and subjected to Western blotting using anti-EP procyclin and anti-GPEET procyclin antibodies (Fig. 5). Substantial decreases in the apparent molecular weights of both the EP and GPEET procyclins harvested from *TbGT10* KO cells, compared to those of wild-type *TbGT10* add-back cells were observed. These data are consistent with a role for TbGT10 in elaborating procyclin GPI anchor sidechains.

**Figure 5.**
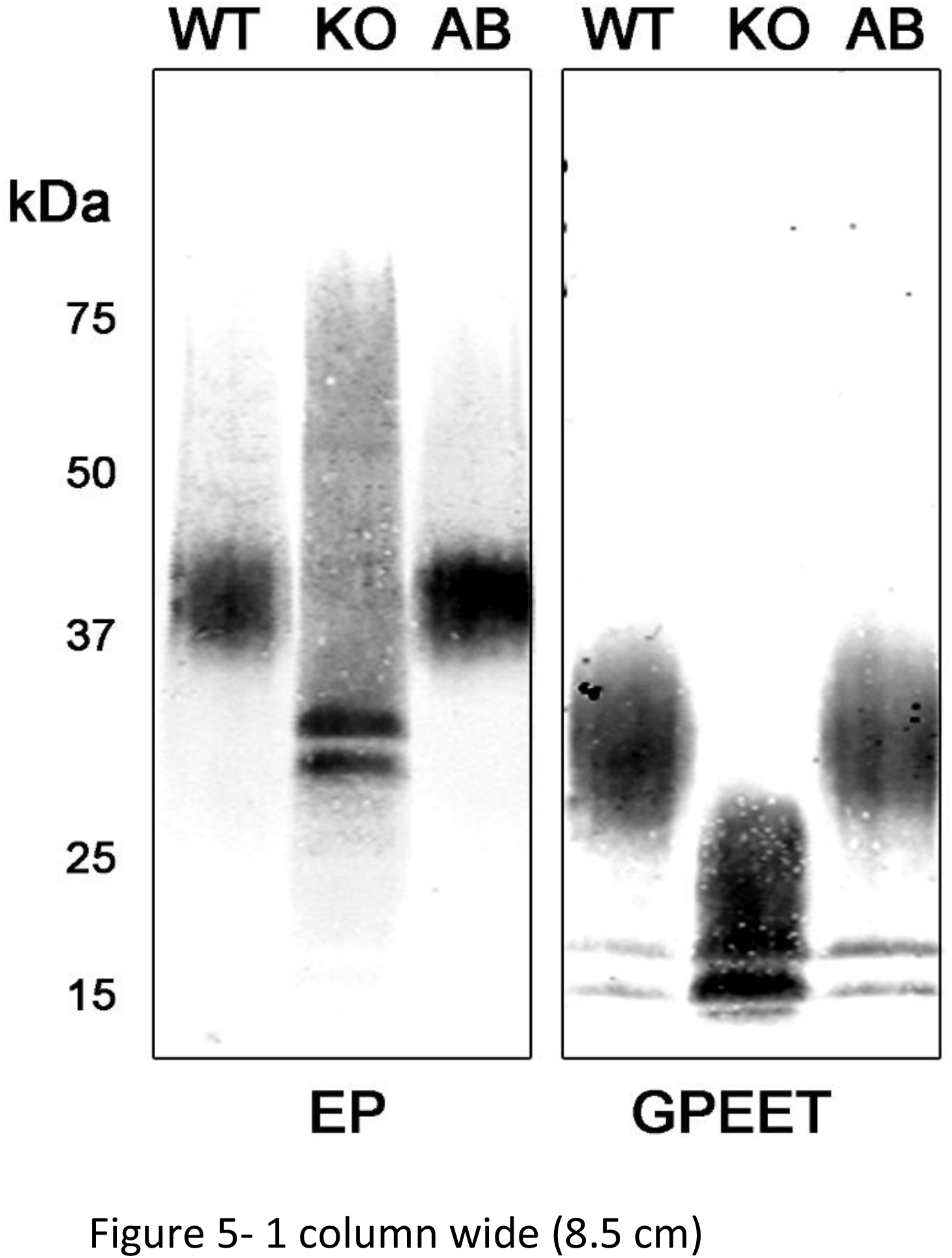
*TbGT10* null procyclic form cells express smaller procyclins than the wild-type. Bloodstream form wild-type (WT), *TbGT10* KO and *TbGT10* AB mutants were differentiated to procyclic form cells *in vitro*. Solvent extracts of each, enriched for procyclins, were resolved by SDS-PAGE, transferred to nitrocellulose by Western blotting and probed with anti-EP and anti-GPEET antibodies as indicated. The positions of molecular weight markers are indicated on the left.

## Discussion

In this paper we describe, for the first time, the application of diCre/*loxP* recombineering (39) in *T. brucei*. We generated cells expressing a single *loxP* flanked *TbGT10* copy at the endogenous locus but found that tetracycline-inducible Cre-recombinase mediated excision was unsuccessful. Therefore, we developed the rapamycin-induced dimerization of Cre approach (38, 45) for use in BSF *T. brucei*. This approach gave efficient conditional (rapamycin-induced) excision of the single *loxP* flanked *TbGT10* copy (*TbGT10*^*Flox*^). This methodology adds another useful approach to the genetic manipulation toolbox for *T. brucei*.

The ablation of TbGT10 had little effect on growth kinetics (Supplementary Fig. S3C) but did produce a fitness cost, evidenced by the outgrowth of cells retaining TbGT10 activity following removal of rapamycin (Supplementary Fig. S4). Limit dilution was used to isolate a stable *TbGT10* KO mutant clone that was viable *in vitro* and able to establish robust murine infections, showing that TbGT10 is non*-* essential to BSF *T. brucei*. Despite this, glycotyping of the BSF *TbGT10* KO mutant revealed striking deficiencies in the synthesis of ricin*-*binding (βGal-terminating) complex *N-*glycans. The ablation of mAb139 and mAbCB1 immunoreactivity in the *TbGT10* KO mutant further suggested an effect on complex *N-*glycosylation. These antibodies detect an previously uncharacterised *N-*linked glycan epitope on the *T. brucei* lysosomal membrane protein (LAMP)-like glycoprotein p67 (Brickman & Balber, 1993; Peck et al., 2008) and on other BSF expressed glycoproteins. Here, using a set of synthetic glycans, we were able to identify the mAb139 and mAbCB1 shared epitope as linear LacNAc repeats of the structure (−<u>6</u>Galβ1-4GlcNAcβ1-)_≥ 4._ These β1-6 interlinked LacNAc repeats are different from the more common β1-3 interlinked (−<u>3</u>Galβ1-4GlcNAcβ1-)_n_ repeats found in mammalian glycoproteins. Methylation linkage analysis of Pronase glycopeptides from BSF wild type and *TbGT10* KO cells showed a reduction in the amount of 3,6-di-*O*-substituted-Gal, representative of -4GlcNAcβ1-6(−4GlcNAcβ1-3)Galβ1-branch points, and of 4-*O*-substituted-GlcNAc and 6-*O*-substituted-Gal representative of -6Galβ1-4GlcNAcβ1-repeats. These data enable us to propose that TbGT10 as a UDP-GlcNAc : βGal β1-6 GlcNAc transferase involved in the synthesis of the large poly-LacNAc *N-*glycan chains unique to BSF *T. brucei* (12). The absence of such glycans has profound effects on the lysosomal processing of p67 and may also be detrimental to p67 activity. We base this on the evidence that p67 knockdown is ultimately lethal, but does not significantly alter lysosomal acidity or endocytic uptake (15). The function of p67 is currently unknown, but RNAi does cause pronounced lysosomal swelling. Interestingly, a recent report gives robust evidence for a possible phospholipase B-like enzymatic or amidase activity (16), whereby the failure to catabolize glycerophospholipids taken up in host serum lipoproteins or inability to fully process degradation substrates results in lysosomal swelling by membrane engorgement or osmotic stress. Our observation that processed p67 peptides accumulate in TbGT10 deficient cells (Fig. 2, KO lane) suggests a similar impairment in lysosomal turnover/export that arises following p67 loss, but crucially we observe a shift in resistance to suramin of a similar magnitude to that manifest by p67 RNAi. However, we do not observe the same dramatic lysosomal swelling or cell death in *TbGT10* KO cells as caused by p67 RNAi, suggesting that p67 function is only partially affected by alterations in its glycosylation as compared to reduction in its polypeptide levels.

Interestingly, removal of TbGT10 was compensated to some extent by an increase 3-*O*-substituted-Gal residues, indicating a concomitant increase in UDP-GlcNAc : βGal β1-3 transferase activity (Fig. 4). The latter is most likely due to TbGT8, a β1-3 GlcNAc-transferase that elaborates PCF procyclin GPI side-chains and BSF complex *N-*glycans (13, 32). As a corollary, we find that BSF *TbGT8* KO glycoproteins react much more strongly with mAb139 (Supplementary Fig. S8). In this case, the absence of -4GlcNAcβ1-6(−4<u>GlcNAcβ1-3</u>)Galβ1-branch points and/or increased TbGT10 activity would be expected to increase the proportion of linear (−6Galβ1-4GlcNAcβ1-)_≥ 4_ motifs and thus mAb139 binding. The outgrowth of TbGT10 expressing cells from the cKO population (Supplementary Fig. S4, lane E) in the absence of rapamycin suggests a fitness cost for impaired glycosylation. In *TbGT10* KO mutants this may compensated by increased TbGT8 activity, whereby the increased synthesis of linear β1-3 interlinked poly-LacNAc chains aids the function of glycoproteins such as p67. A *TbGT10*/*TbGT8* double null mutant, if viable, would be a useful model to explore such a hypothesis.

The detection of low apparent MW procyclins in our PCF TbGT10 KO mutant implicates TbGT10 as a bifunctional enzyme, working not only on BSF complex *N*-glycans but also on procyclin GPI sidechains. In Fig. 6 we provide models of how we think ablation of TbGT10 (this paper), TbGT8 (13, 32) and TbGT3 (31) affect BSF complex *N*-link structures and procyclin GPI sidechain structures.

In summary, we propose that TbGT10 is a bifunctional UDP-GlcNAc : βGal β1-6 GlcNAc-transferase active in both BSF and PCF life-cycle stages elaborating complex *N*-glycans and GPI sidechains, respectively. This provides further evidence that *T. brucei* has evolved a large UDP-GlcNAc/UDP-Gal-dependent glycosyltransferase repertoire from an ancestral β3GT gene, and that members of this family can catalyse the formation of the β1-2, β1-3 and (this study) β1-6 glycosidic linkages found in its glycoproteins.

**Figure 6.**
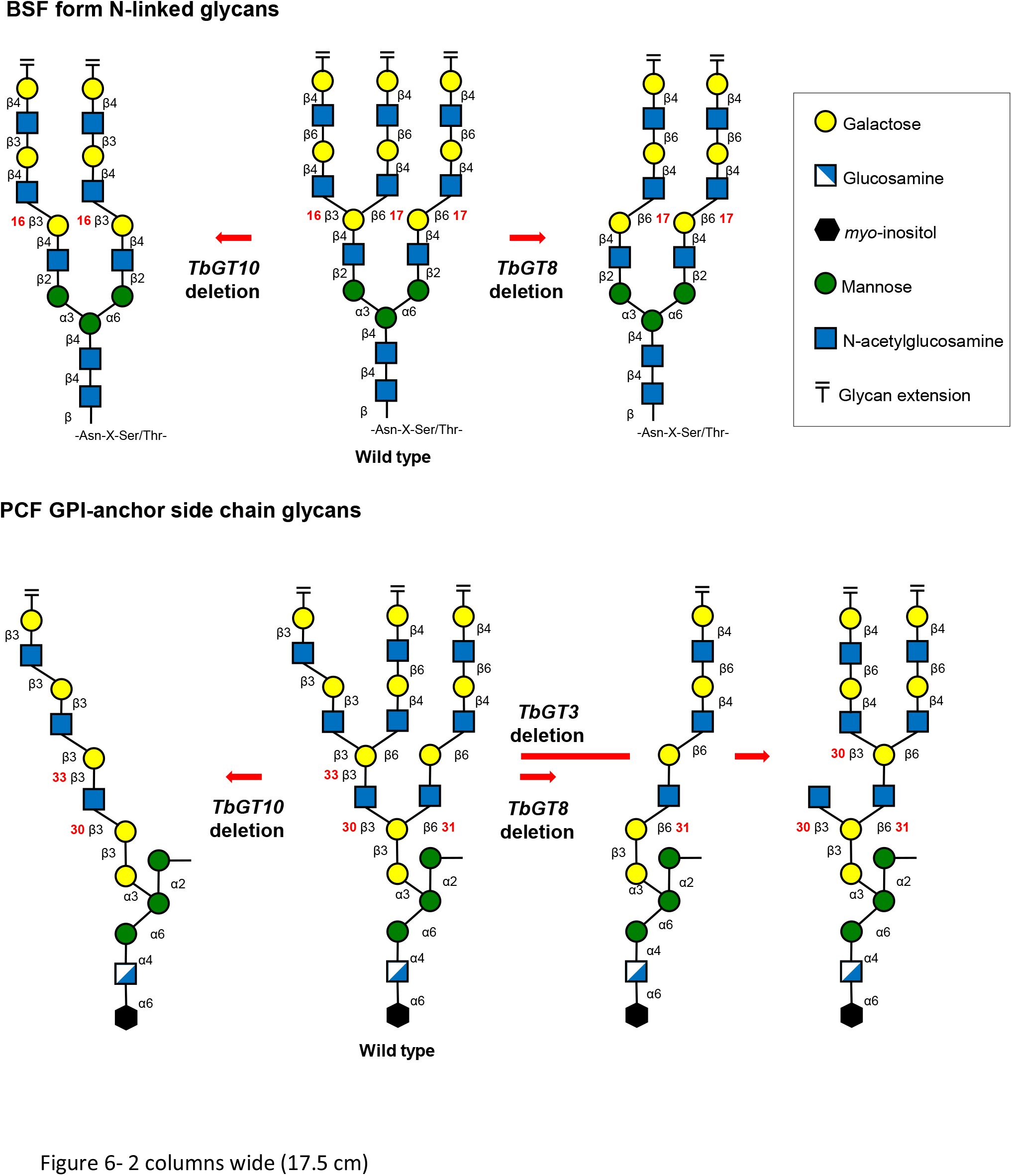
Current model of *N*-linked and GPI anchor glycan biosynthesis in *T. brucei* by analysis of TbGT mutants. The glycolinkages catalysed by experimentally characterised TbGTs are represented by red numbers (see Supplementary Table S1). The loss of UDP-GlcNAc : βGal transferase activity due to TbGT8 and TbGT10 deletion prevents glycolinkages 16 and 17, respectively, resulting in deficiencies in the synthesis of 4GlcNAcβ1-3(−4GlcNAcβ1-6)Galβ1-branch points in *N*-linked glycans. TbGT8 and TbGT10 are bifunctional and also form glycolinkages 30 and 31 required for GPI glycan side-chain synthesis. Loss of either glycosyltransferase activity similarly prevents branch point formation and results in the expression of low apparent MW procyclins. Deletion of *TbGT3* ablates UDP-Gal : βGlcNAc β1-3 Gal transferase activity (linkage 33) on the 3-arm of the GPI-anchor side chain. Elaboration and branch point synthesis occurs on the 6-arm due to TbGT8/10 activity and an unidentified UDP-Gal : βGlcNAc β1-4 GalT.

## Acknowledgements

We thank George Cross for making *loxP* constructs accessible, James Bangs for mAb139, mAbCB1 and mAbp67C and for helpful discussions, David Robinson for assistance with the BLI assay, and Lucia Guther for valuable experimental advice and the suggestion to look at the effects of *TbGT10* knockout on p67. This work was supported by a Wellcome Trust Investigator Award to MAJF (101842/Z13/Z).

## MATERIALS AND METHODS

### Cultivation of Trypanosomes

*Trypanosoma brucei brucei* Lister strain 427 bloodstream form parasites, expressing VSG variant 221 (MiTat1.2) and transformed to stably express T7 polymerase and the tetracycline repressor protein under G418 antibiotic selection, were used in this study. This genetic background will be referred to from hereon as wild-type (WT). Cells were cultivated in HMI-11 medium containing 2.5 μg/ml G418 at 37 °C in a 5% CO_2_ incubator as described in (46). Differentiation to procyclic-form *T. brucei* cells was initiated by washing ∼ 5 × 10^8^ of log stage bloodstream-form cells in SDM-79 medium then culturing in 500 mL of SDM-79 media supplemented with 15 % FBS and 3 mM *cis*-aconitate at 28°C. Cells were harvested after 4 days culture.

### DNA Isolation and Manipulation

Plasmid DNA was purified from *Escherichia coli* DH5α competent cells (MRC PPU Reagents & Services, Dundee) using a Qiagen Miniprep kit. Gel extraction and reaction clean-up was performed using Qiaquick kits (Qiagen). Custom oligonucleotides were obtained from Thermo Fisher. *T. brucei* genomic DNA was isolated from ∼ 2 × 10^8^ bloodstream form cells using a DNeasy Blood & Tissue Kit (Qiagen) using standard methods.

### Generation of Gene Replacement Constructs

A full list and descriptions of all primers (Table S2) used in this study are available. The puromycin acetyltransferase (*PAC*) drug resistance cassette was generated by PCR-amplification of 5’-537 bp and 3’-530 bp flanking regions of Tb427.5.2760 using *Pfu* DNA polymerase with primers JR1/2 and JR3/4 respectively. The two PCR products were used together in a further PCR reaction to yield a product containing the 5’ flank linked to the 3’ flank by a short *HindIII, PmeI* and *BamHI* cloning site. The *PAC* drug-resistance gene was then introduced into the targeting vector via the *HindIII* and *BamHI* cloning sites. The first allele of *GT10* was replaced with the *PAC* drug resistance construct to generate a Δ*gt10*::*PAC*/*GT10* single deletion mutant. The *loxP* expression construct containing a floxed MCS conferring a C-terminal 3xHA epitope tag in tandem array with *HYG_TK* selectable markers (*pSY45_pDS66*) was generated by restriction digestion of pSY45 and pDS66 constructs with *BglIII* and *Nde*I and subsequent T4 Ligation. Q5 polymerase PCR amplification of Tb427.5.2760 lacking a 3’ stop codon using oligonucleotides SMD169/186 to confer 5’ *FseI* and 3’ *BglIII* cloning sites for cloning into the *loxP* vector to generate *pSY45_pDS66_GT10cHA* _*3*_ ^*Flox*^. This construct was used as a template for PCR amplification using SMD173/4 whilst 5’-535 bp and 3’ 485 bp homologous flanks for Tb927.5.2760 were amplified using SMD171/2 and SMD175/176, respectively. A linear pUC19 acceptor vector was generated using Q5 polymerase and primers SMD33/4. Each PCR amplicon contained 20 bp overlapping ends to facilitate Gibson assembly. Amplicons were PCR purified (Qiagen) and 10 pmol of each used in a 4-fragment Gibson assembly reaction (NEB) to generate *GT10 5’_pSY45_pDS66_GT10cHA*_*3*_ ^*Flox*^*_GT10 3’_pUC19*. The vector was PCR amplified using NEBase Changer and mutagenesis primers SMD181/2 introducing a stop codon (TAG) at the 3’ end of the *TbGT10* ORF to prevent epitope tagging. This construct was used to generate the Δ*gt10*::*PAC*/Δ*gt10*::*GT10*^*FLox*,^ cell line hereon referred to as ‘*GT10 flox’*, and later used to generate Δ*gt10*::*PAC*/Δ*gt10*::*GT10*^*FLox*^ (*SSU diCre*) conditional knockout mutants (cKO). Stable null mutants lacking *GT10*^*Flox*^ following diCre mediated excision (KO) were obtained with this cell line by limit dilution and further complemented by re-integration of this construct to generate functional *TbGT10* add-back (AB) mutants. To generate a diCre expression vector, the *Cre59-FKBP12* and *Cre60-FRB* coding sequences separated by a 330 bp *TbActin* intergenic regulatory element were assembled *in silico* using CLC Main Workbench software (Qiagen). The sequence was synthesised (Dundee Cell Products) and cloned into pLEW100v5 by *XhoI* and *BamHI* restriction enzyme sites. A 92bp region containing the Tet repressor operons were deleted by Q5 polymerase mediated mutagenesis using oligonucleotides SMD277/8 to confer constitutive expression. The identity of all constructs was confirmed by sequencing.

### Transformation of bloodstream form T. brucei

Constructs for gene replacement and ectopic expression were purified, digested with appropriate restriction enzymes to linearize, precipitated, washed with 70% ethanol, and re-dissolved in sterile water. The linearized DNA was electroporated into *T. brucei* bloodstream form cells (Lister strain 427, variant 221) that were stably transformed to express T7 RNA polymerase and the tetracycline repressor protein under G418 selection. Cell transformation was carried out as described previously (46–48).

### Induction of diCre mediated gene deletion

Mid-log stage *TbGT10* cKO cultures (∼1 × 10^6^ cells/mL) were passaged to 2 × 10^3^ cells/mL or 2 × 10^4^ cells/mL and dosed with 100 nM rapamycin (Abcam) from a 1 mM stock solution in DMSO. Cells at late-log phase were harvested for analysis after 3 or 2 days respectively. Conditional gene deletion was assessed by PCR amplification of genomic DNA using Taq polymerase (NEB) and oligonucleotides SMD157/160 flanking the *TbGT10* locus. Hygromycin drug sensitivity was used as a proxy of *TbGT10*^*Flox*^ loss by passaging cells in the presence or absence of 100 nM rapamycin for 3 days before seeding at 2x 10^3^ cells/mL and culturing for 3 days in the presence of hygromycin. Daily cell counting was performed to assess growth rates. A single clone that gave a robust growth defect in the presence of hygromcyin was selected as the *TbGT10* cKO mutant.

### Generation of TbGT10 KO mutants

*TbGT10* cKO cells grown in the presence of 100 nM rapamycin for 5 days were cultured in the presence of 100 µM ganciclovir (GCV) for a further 3 days. Cells were serially diluted in 96 well plates in the presence of 100 µM GCV and 100 nM rapamycin. Four clones were seeded at 2 × 10^3^ cells/mL +/- hygromycin to confirm loss of *TbGT10_TK-HYG*^*Flox*^ array by drug sensitivity. TbGT10 loss was confirmed by mAb139 immunodetection and a single clone lacking *TbGT10* (KO) was selected for mouse infection, complementation and GC-MS linkage analysis.

### Mouse infectivity studies

Wild-type, *TbGT10* KO and *TbGT10* add-back (AB) mutant bloodstream form trypanosomes were grown in HMI-11T media, washed in media without antibiotics and re-suspended at 1 × 10^6^ cells/ml. Groups of 6 female Balb/c mice were used for each cell line and 0.2 ml of the parasite suspension was injected intraperitoneally per animal. Infections were assessed 2 and 3 days post-infection by tail bleeding and cell counting using a Neubauer chamber in a phase contrast microscope. At day 3 post-infection, parasites were pooled from the blood of 3 mice in the same treatment group and BSF genomic DNA was extracted. *TbGT10* KO was confirmed by performing PCR amplification with oligonucleotides SMD111/2 alongside a control gene (TbGTZ) using SMD109/110.

### Western/ Lectin blotting

For Western and lectin blot analysis, 5 × 10^6^ – 1 × 10^7^ cells were lysed in cells were lysed in 25mM Tris, pH 7.5, 100 mM NaCl, 1% Triton X-100 and solubilised in 1xSDS sample buffer containing 0.1 M DTT by heating at 55°C for 20 mins. Glycoproteins were resolved by SDS–PAGE (approx. 1×10^7^ cell equivalents/lane) on NuPAGE *bis*-Tris 4–12% gradient acrylamide gels (Invitrogen) and transferred to nitrocellulose membrane (Invitrogen). Ponceau S staining confirmed equal loading and transfer. Glycoproteins were probed with 1.7 μg/ml biotin-conjugated ricin (RCA-120, Vector Laboratories, UK) in blocking buffer before or after pre-incubation with 10 mg/ml D-galactose and 10 mg/ml lactose to confirm specific ricin binding. Detection was performed using IRDye 680LT-conjugated Streptavidin and the LI-COR Odyssey Infrared Imaging System (LICOR Biosciences). For Western blotting, monoclonal antibodies mAb139 and CB1 and polyclonal p67C were diluted 1:1000 in blocking buffer (50mM Tris-HCl pH7.4, 0.15M NaCl, 0.25% BSA, 0.05% (w/v) Tween-20, 0.05% NaN_3_ and 2% (w/v) Fish Skin Gelatin). Detection with IRDye® 800CW Goat anti-Mouse and anti-Rabbit (LICOR Biosciences) was performed using 1:15,000 dilutions in blocking buffer. Procyclins purified by butan-1-ol phase separation (49) from 1 × 10^7^ cell equivalents were resolved by 10% SDS-PAGE gel electrophoresis and transferred to nitrocellulose membranes. Western blotting with monoclonal anti-EP and polyclonal anti-GPEET was performed at 1:750 dilutions and detected using IRDye® 800CW Goat anti-Mouse and IRDye® 680RD Donkey anti-Rabbit at 1:15,000 and 1:20,000 respectively.

### EC50 assay

Wild-type cells were used to assess rapamycin sensitivity and *TbGT10* cKO cells were pre-induced for 3 days in the presence or absence of 100 nM rapamycin to assess suramin sensitivity. For EC50 determination, cells were seeded at 2 x10^3^ cells/mL in 96 well plates in a rapamycin or suramin 2-fold dilution series. Tetracycline-induced and uninduced cells were grown in triplicate rows of the same plate. After 66 h growth, 20 uL of 0.125 mg/ mL resazurin in HMI-11 was added to each well and the plates incubated for a further 6 h. Fluorescence was measured using a Tecan plate reader at an excitation wavelength of 528 nM and emission wavelength of 590 nM. Data were processed by using GRAFIT (version 5.0.4; Erithacus Software) and fitted to a 2-parameter equation, where the data are corrected for background fluorescence, to obtain the effective concentration inhibiting growth by 50% (EC50).

### Bio-layer interferometry analysis

Synthetic biotinylated LacNAc oligosacharides were synthesised as described in (42) and (Supplementary Fig. 6). Biolayer interferometry (BLI) measurements were carried out using an Octet RED 384 instrument (ForteBio). Each biotinylated LacNAc oligosaccharide (15 nM solution in 1 x phosphate buffered saline) was immobilised by incubation for 10 min with superstreptavidin (SSA) biosensor pins, alongside a buffer only control. Any unbound streptavidin-binding sites were blocked by a 1 min dip into 10 mg/mL biocytin (Tocris). Each set of biosensors was incubated with 25 nM solutions of IgG2 mAb139 in 1 x phosphate buffered saline in 6 min association/disassociation cycles. Regeneration was performed by eluting bound mAb139 by incubation in 0.2M glycine-HCl, pH 2.8, and quenching in 1 M Tris-HCl (pH 7.0) before repeating the association/disassociation cycle twice more. After the final 0.2M glycine-HCl, pH 2.8, incubation the assay was repeated using 25 nM IgG as an isotype control and the association/disassociation/regeneration cycle repeated thrice. Sensors were washed for 30 s in 1 x phosphate buffered saline between each step except for the association and disassociations. The single step assay was performed as described but using 15 nM of IgM CB1 or mAb139 with 5 min association and 15 min disassociation steps.

### N-glycopeptide preparation for GC-MS analysis

To prepare *N*-glycopeptides of *T. brucei* bloodstream-form wild-type and *TbGT10* KO mutant cells, 5 × 10^9^ cells were harvested and washed twice with 1X trypanosome dilution buffer (TDB; 5 mM KCl, 80 mM NaCl, 1 mM MgSO_4_, 20 mM Na_2_ HPO_4_, 2 mM NaH_2_ PO_4_, 20 mM glucose, pH 7.4) and depleted of sVSG using hypotonic lysis. Briefly, the cells were lysed by osmotic shock by incubating with water containing 0.1 mM TLCK, 1 µg/ml leupeptin and 1 µg/ml aprotinin (pre-warmed to 37°C) at 37 °C for 5 min. This releases all the cystosolic components and majority of VSG protein as soluble form VSG (sVSG). The sVSG depleted cell ghost pellet was resuspended in 1.5 ml of 20 mM ammonium bicarbonate and mixed with 50 µl of 10 mg/ml of freshly prepared Pronase (Calbiochem, #53702) dissolved in 5 mM calcium acetate and digested at 37 °C for 24 h. The digest was centrifuged at 12000 x g for 30 min to remove the cell ghost membranes and nuclei and the supernatant was incubated at 95 °C for 20 min to heat inactivate the Pronase and again centrifuged to remove particulates. The supernatant containing the Pronase digested glycopeptides was applied to a 30 kDa cut-off centrifugal filter (Amicon) and diafiltrated with water three times. The resulting aqueous fraction was subjected to chloroform phase separation by mixing with equal volume of chloroform to remove any remaining lipid contaminants. The upper aqueous phase (Pronase glycopeptide fraction) was collected in a fresh tube and used for GC-MS monosaccharide composition and methylation linkage analysis.

### Monosaccharide composition and methylation linkage analysis

Aliquots (10%) of the Pronase glycopeptide fractions were dried and mixed with 1 nmol *scyllo*-inositol internal standard and subjected to methanolysis, re-*N*-acetylation and trimethylsilylation and GC-MS monosaccharide composition analysis of the resulting 1-*O*-methyl-glycoside TMS derivatives, as described in (49). Remaining 90% of the samples were used for methylation linkage analysis (49). Briefly, the samples were dried and subjected to permethylation using the sodium hydroxide method. The permethylated glycans were then subjected to acid hydrolysis, NaB(^2^H)_4_ reduction, and acetylation to generate partially methylated alditol acetates (PMAAs). The 1-*O*-methyl-glycoside TMS derivatives and the PMAAs were analysed by GC-MS (Agilent Technologies, 7890B Gas Chromatography system with 5977A MSD, equipped with Agilent HP-5ms GC Column, 30 m X 0.25 mm, 0.25 µm) as described in (49).

## References

1. Capewell, P., Cren-Travaillé, C., Marchesi, F., Johnston, P., Clucas, C., Benson, R. A., Gorman, T.-A., Calvo-Alvarez, E., Crouzols, A., Jouvion, G., Jamonneau, V., Weir, W., Stevenson, M. L., O’Neill, K., Cooper, A., Swar, N. K., Bucheton, B., Ngoyi, D. M., Garside, P., Rotureau, B., and MacLeod, A. (2016) The skin is a significant but overlooked anatomical reservoir for vector-borne African trypanosomes. Elife. 10.7554/eLife.17716

2. Trindade, S., Rijo-Ferreira, F., Carvalho, T., Pinto-Neves, D., Guegan, F., Aresta-Branco, F., Bento, F., Young, S. A., Pinto, A., Van Den Abbeele, J., Ribeiro, R. M., Dias, S., Smith, T. K., and Figueiredo, L. M. (2016) Trypanosoma brucei Parasites Occupy and Functionally Adapt to the Adipose Tissue in Mice. Cell Host Microbe. 19, 837–848

3. Cross, G. A. M. (1996) Antigenic variation in trypansosomes: Secrets surface slowly. BioEssays. 18, 283–291

4. Mehlert, A., Bond, C. S., and Ferguson, M. A. J. J. (2002) The glycoforms of a Trypanosoma brucei variant surface glycoprotein and molecular modeling of a glycosylated surface coat. Glycobiology. 12, 607–12

5. Mehlert, A., Zitzmann, N., Richardson, J. M., Treumann, A., and Ferguson, M. A. J. (1998) The glycosylation of the variant surface glycoproteins and procyclic acidic repetitive proteins of Trypanosoma brucei. Mol. Biochem. Parasitol. 91, 145–152

6. Schwede, A., Macleod, O. J. S., MacGregor, P., and Carrington, M. (2015) How Does the VSG Coat of Bloodstream Form African Trypanosomes Interact with External Proteins? PLoS Pathog. 11, 1–18

7. Hartel, A. J. W., Glogger, M., Jones, N. G., Abuillan, W., Batram, C., Hermann, A., Fenz, S. F., Tanaka, M., and Engstler, M. (2016) N-glycosylation enables high lateral mobility of GPI-anchored proteins at a molecular crowding threshold. Nat. Commun. 7, 12870

8. Horn, D. (2014) Antigenic variation in African trypanosomes. Mol. Biochem. Parasitol. 195, 123–9

9. Pinger, J., Nešić, D., Ali, L., Aresta-Branco, F., Lilic, M., Chowdhury, S., Kim, H.-S., Verdi, J., Raper, J., Ferguson, M. A. J., Papavasiliou, F. N., and Stebbins, C. E. (2018) African trypanosomes evade immune clearance by O-glycosylation of the VSG surface coat. Nat. Microbiol. 3, 932–938

10. Zamze, S. E., Wooten, W., Ashford, D. A., Ferguson, M. A. J., Dwek, R. A., Rademacher, T. W., and Zamze, S. E. (1990) Characterisation of the asparagine-linked oligosaccharides from Trypanosoma brucei type-I variant surface glycoproteins

11. Zamze, S. E., Ashford, D. A., Wooten, E. W., Rademacher, T. W., and Dwek, R. A. (1991) Structural characterization of the asparagine-linked oligosaccharides from Trypanosoma brucei Type II and Type III variant surface glycoproteins. J. Biol. Chem. 266, 20244–20261

12. Atrih, A., Richardson, J. M., Prescott, A. R., and Ferguson, M. A. J. (2005) Trypanosoma brucei Glycoproteins Contain Novel Giant Poly-N-acetyllactosamine Carbohydrate Chains. J. Biol. Chem. 280, 865–871

13. Izquierdo, L., Nakanishi, M., Mehlert, A., Machray, G., Barton, G. J., and Ferguson, M. a J. (2009) Identification of a glycosylphosphatidylinositol anchor-modifying β1-3 N-acetylglucosaminyl transferase in Trypanosoma brucei. Mol. Microbiol. 71, 478–491

14. Alexander, D. L., Schwartz, K. J., Balber, A. E., and Bangs, J. D. (2002) Developmentally regulated trafficking of the lysosomal membrane protein p67 in Trypanosoma brucei. J. Cell Sci. 115, 3253–63

15. Peck, R. F., Shiflett, A. M., Schwartz, K. J., McCann, A., Hajduk, S. L., and Bangs, J. D. (2008) The LAMP-like protein p67 plays an essential role in the lysosome of African trypanosomes. Mol. Microbiol. 68, 933–946

16. Koeller, C. M., Smith, T. K., Gulick, A. M., and Bangs, J. D. (2020) P67: A cryptic lysosomal hydrolase in Trypanosoma brucei ? Parasitology. 10.1017/S003118202000195X

17. Nolan, D. P., Geuskens, M., and Pays, E. (1999) N-linked glycans containing linear poly-N-acetyllactosamine as sorting signals in endocytosis in Trypanosoma brucei. Curr. Biol. 10.1016/S0960-9822(00)80018-4

18. Roditi, I., Schwarz, H., Pearson, T. W., Beecroft, R. P., Liu, M. K., Richardson, J. P., Buhring, H. J., Pleiss, J., Bulow, R., Williams, R. O., and Overath, P. (1989) Procyclin gene expression and loss of the variant surface glycoprotein during differentiation of Trypanosoma brucei. J. Cell Biol. 108, 737–746

19. Treumann, A., Zitzmann, N., Hülsmeier, A., Prescott, A. R., Almond, A., Sheehan, J., and Ferguson, M. A.. (1997) Structural characterisation of two forms of procyclic acidic repetitive protein expressed by procyclic forms of Trypanosoma brucei. J. Mol. Biol. 269, 529–547

20. Mehlert, A., Treumann, A., and Ferguson, M. A. J. (1999) Trypanosoma brucei GPEET-PARP is phosphorylated on six out of seven threonine residues. Mol. Biochem. Parasitol. 98, 291–296

21. Ferguson, M. A. J., Murray, P., Rutherford, H., and Mcconville, M. J. (1993) A simple purification of procyclic acidic repetitive protein and demonstration of a sialylated glycosyl-phosphatidylinositol membrane anchor, [online]

22. Engstler, M., Reuter, G., and Schauer, R. (1993) The developmentally regulated trans-sialidase from Trypanosoma brucei sialylates the procyclic acidic repetitive protein. Mol. Biochem. Parasitol. 61, 1–13

23. Nagamune, K., Acosta-Serrano, A., Uemura, H., Brun, R., Kunz-Renggli, C., Maeda, Y., Ferguson, M. A. J., and Kinoshita, T. (2004) Surface Sialic Acids Taken from the Host Allow Trypanosome Survival in Tsetse Fly Vectors. J. Exp. Med. 199, 1445–1450

24. Acosta-Serrano, A., Vassella, E., Liniger, M., Kunz Renggli, C., Brun, R., Roditi, I., and Englund, P. T. (2001) The surface coat of procyclic Trypanosoma brucei: Programmed expression and proteolytic cleavage of procyclin in the tsetse fly. Proc. Natl. Acad. Sci. 98, 1513–1518

25. Izquierdo, L., Schulz, B. L., Rodrigues, J. A., Güther, M. L. S., Procter, J. B., Barton, G. J., Aebi, M., and Ferguson, M. A. J. (2009) Distinct donor and acceptor specificities of Trypanosoma brucei oligosaccharyltransferases. EMBO J. 28, 2650–2661

26. Izquierdo, L., Mehlert, A., and Ferguson, M. A. (2012) The lipid-linked oligosaccharide donor specificities of Trypanosoma brucei oligosaccharyltransferases. Glycobiology. 22, 696–703

27. Jinnelov, A., Ali, L., Tinti, M., Lucia, M., Güther, S., Michael, X., and Ferguson, A. J. (2017) Single-subunit oligosaccharyltransferases of Trypanosoma brucei display different and predictable peptide acceptor specificities. 10.1074/jbc.M117.810945

28. Poljak, K., Breitling, J., Gauss, R., Rugarabamu, G., Pellanda, M., and Aebi, M. (2017) Analysis of substrate specificity of Trypanosoma brucei oligosaccharyltransferases (OSTs) by functional expression of domain-swapped chimeras in yeast. J. Biol. Chem. 292, 20342–20352

29. Cantarel, B. L., Coutinho, P. M., Rancurel, C., Bernard, T., Lombard, V., and Henrissat, B. (2009) The Carbohydrate-Active EnZymes database (CAZy): an expert resource for Glycogenomics. Nucleic Acids Res. 37, 233–238

30. Narimatsu, H. (2006) Human glycogene cloning: focus on β3-glycosyltransferase and β4-glycosyltransferase families. Curr. Opin. Struct. Biol. 16, 567–575

31. Izquierdo, L., Acosta-Serrano, A., Mehlert, A., and Ferguson, M. A. (2015) Identification of a glycosylphosphatidylinositol anchor-modifying β1-3 galactosyltransferase in Trypanosoma brucei. Glycobiology. 25, 438–47

32. Nakanishi, M., Karasudani, M., Shiraishi, T., Hashida, K., Hino, M., Ferguson, M. A. J., and Nomoto, H. (2014) TbGT8 is a bifunctional glycosyltransferase that elaborates N-linked glycans on a protein phosphatase AcP115 and a GPI-anchor modifying glycan in Trypanosoma brucei. Parasitol. Int. 63, 513–8

33. Damerow, M., Rodrigues, J. a, Wu, D., Güther, M. L. S., Mehlert, A., and Ferguson, M. a J. (2014) Identification and Functional Characterization of a Highly Divergent N-Acetylglucosaminyltransferase I (TbGnTI) in Trypanosoma brucei. J. Biol. Chem. 289, 9328–39

34. Damerow, M., Graalfs, F., Güther, M. L. S., Mehlert, A., Izquierdo, L., and Ferguson, M. A. J. (2016) A Gene of the β3-Glycosyltransferase Family Encodes N -Acetylglucosaminyltransferase II Function in Trypanosoma brucei. J. Biol. Chem. 291, 13834–13845

35. Alsford, S., Eckert, S., Baker, N., Glover, L., Sanchez-Flores, A., Leung, K. F., Turner, D. J., Field, M. C., Berriman, M., and Horn, D. (2012) High-throughput decoding of antitrypanosomal drug efficacy and resistance. Nature. 482, 232–236

36. Wiggins, C. A. R., and Munro, S. (1998) Activity of the yeast MNN1 α-1,3-mannosyltransferase requires a motif conserved in many other families of glycosyltransferases. Proc. Natl. Acad. Sci. U. S. A. 95, 7945–7950

37. Jullien, N., Sampieri, F., Enjalbert, A., and Herman, J. (2003) Regulation of Cre recombinase by ligand-induced complementation of inactive fragments. Nucleic Acids Res. 31, e131

38. Duncan, S. M., Myburgh, E., Philipon, C., Brown, E., Meissner, M., Brewer, J., and Mottram, J. C. (2016) Conditional gene deletion with DiCre demonstrates an essential role for CRK3 in Leishmania mexicana cell cycle regulation. Mol. Microbiol. 100, 931–44

39. Kim, H. S., Li, Z., Boothroyd, C., and Cross, G. a M. (2013) Strategies to construct null and conditional null Trypanosoma brucei mutants using Cre-recombinase and loxP. Mol. Biochem. Parasitol. 191, 16–19

40. Barquilla, A., Crespo, J. L., and Navarro, M. (2008) Rapamycin inhibits trypanosome cell growth by preventing TOR complex 2 formation. Proc. Natl. Acad. Sci. U. S. A. 105, 14579–14584

41. Brickman, M. J., and Balber, A. E. (1993) Trypanosoma brucei rhodesiense: Membrane Glycoproteins Localized Primarily in Endosomes and Lysosomes of Bloodstream Forms. Exp. Parasitol. 76, 329–344

42. Buzzi, B. (2008) Synthesis of oligosaccharides and glycoconjugates as antigens for vaccine formulation. Ph.D. thesis, University of Milan, Faculty of Mathematical, Physical and Natural Sciences, Doctoral School in Chemical Sciences and Technologies

43. Ferguson, M. A. J., Low, M. G., and Cross, G. A. M. (1985) Glycosyl-sn-1,2-dimyristylphosphatidylinositol is covalently linked to Trypanosoma brucei variant surface glycoprotein. J. Biol. Chem. 260, 14547–14555

44. De Almeida, M. L. C., and Turner, M. J. (1983) The membrane form of variant surface glycoproteins of Trypanosoma brucei. Nature. 302, 349–352

45. Kangussu-Marcolino, M. M., Cunha, A. P., Avila, A. R., Herman, J.-P., and DaRocha, W. D. (2014) Conditional removal of selectable markers in Trypanosoma cruzi using a site-specific recombination tool: Proof of concept. Mol. Biochem. Parasitol. 198, 71–74

46. Wirtz, E., Leal, S., Ochatt, C., and Cross, G. A. M. (1999) A tightly regulated inducible expression system for conditional gene knock-outs and dominant-negative genetics in Trypanosoma brucei. Mol. Biochem. Parasitol. 99, 89–101

47. Güther, M. L. S., Leal, S., Morrice, N. A., Cross, G. A. M., and Ferguson, M. A. J. (2001) Purification, cloning and characterization of a GPI inositol deacylase from Trypanosoma brucei. EMBO J. 20, 4923–4934

48. Milne, K. G., Güther, M. L. S., and Ferguson, M. A. J. (2001) Acyl-CoA binding protein is essential in bloodstream form Trypanosoma brucei. Mol. Biochem. Parasitol. 112, 301–304

49. Ferguson, M. A. J. (1993) GPI membrane anchors: isolation and analysis. in Glycobiology (Fukuda, M., and Kobata, A. eds), pp. 349–383, Oxford: IRL Press at Oxford University Press

